# Maternal prenatal distress exposure negatively associates with the stability of neonatal frontoparietal network

**DOI:** 10.1101/2023.01.17.524480

**Authors:** Jetro J. Tuulari, Olli Rajasilta, Joana Cabral, Morten L. Kringelbach, Linnea Karlsson, Hasse Karlsson

**Affiliations:** FinnBrain Birth Cohort Study, Turku Brain and Mind Center, Department of Clinical Medicine, University of Turku, Turku, Finland; Department of Clinical Medicine, Psychiatry, University of Turku and Turku University Hospital, Turku, Finland; Centre for Eudaimonia and Human Flourishing, Linacre College, University of Oxford, Oxford, UK; Department of Psychiatry, University of Oxford, Oxford, UK; Life and Health Sciences Research Institute (ICVS), School of Medicine, University of Minho, Braga, Portugal; ICVS/3B’s—PT Government Associate Laboratory, Braga/Guimarães, Portugal; Turku Collegium for Science, Medicine and Technology (TCSMT), University of Turku, Turku, Finland; Centre for Population Health Research, University of Turku and Turku University Hospital, Turku Finland; Department of Clinical Medicine, Paediatrics and Adolescent Medicine, University of Turku and Turku University Hospital, Turku, Finland; Centre for Music in the Brain, Aarhus University, Aarhus, Denmark

## Abstract

Maternal prenatal distress (PD), frequently defined as *in utero* prenatal stress exposure (PSE) to the developing fetus, influences the developing brain and numerous associations between PSE and brain structure have been described both in neonates and in older children. Previous studies addressing PSE-linked alterations in neonates’ brain activity have focused on connectivity analyses from predefined seed regions, but the effects of PSE at the level of distributed functional networks remains unclear. In this study, we investigated the impact of prenatal distress on the spatial and temporal properties of functional networks detected in functional MRI data from 20 naturally sleeping, term-born (age 25.85 ± 7.72 days, 11 males), healthy neonates. First, we performed group level independent component analysis (GICA) to evaluate an association between PD and the spatial configuration of the functional networks. Second, we search for an association with PD at the level of the stability of functional networks over time using leading eigenvector dynamics analysis (LEiDA). No statistically significant associations were detected at the spatial level for the GICA-derived networks. However, at the dynamic level, LEiDA revealed that maternal PD significantly decreased the stability of a frontoparietal network. These results imply that maternal PD may influence the stability of frontoparietal connections in neonatal brain network dynamics and adds to the cumulating evidence that frontal areas are especially sensitive to PSE.

## Introduction

Maternal prenatal distress (PD) is common as ca. 30% of pregnant women report psychosocial stress via work-related stress as well as depressive / anxiety symptoms (1). This may be relevant for transmission of intergenerational health risks (2). Correspondingly, maternal PD can be conceptualised as *in utero* prenatal stress exposure (PSE) to the fetus. PSE has potentially numerous adverse effects on later development such as increased risk for psychiatric disorders (3) and less advanced cognitive development (2), including numerous phenotypes that are closely linked to brain function. Indeed, numerous studies have indicated that there are close links between PSE and child’s structural brain development (4).

Studies addressing PSE-linked functional associations have been scarce and mainly used seed-based connectivity analyses of the bilateral amygdala (5–8). In these studies, the functional connectivity from amygdala to subcortical and prefrontal regions had negative associations with different measures of PSE, mainly maternal depression and anxiety symptoms, quantified by a binary system of maternal clinical diagnosis of depression/anxiety (5), Center for Epidemiological Studies Depression Scale (CES-D) (6) and Edinburgh Postnatal Depression Scale (EPDS) (7,8). Our recent work extended the analyses to whole brain voxel-wise activity profiles and this work implicated a positive association between neonatal fractional amplitude of low-frequency fluctuations (fALFF) in ventromedial prefrontal cortex (vmPFC) and a positive association of frontoparietal areas with vmPFC seed-based connectivity (9). No studies have however, mapped the effects of PSE to neonate distributed functional networks, both in terms of their spatial configuration or their dynamics over time.

In the current study, we used a composite score of maternal depressive and anxiety symptoms as a measure for PSE. We analyzed functional MRI data from 20 naturally sleeping, term-born (gestational age 43.7, SD 0.7 weeks) healthy neonates acquired over 6 minutes with a combination of independent component analysis (ICA) and a novel method to evaluate the stability of functional networks over time, called Leading Eigenvector Dynamics Analysis (LEiDA) (10). We hypothesized, as observed in our prior study (9), that PSE would be associated with the dynamics of these networks. This study was explorative and thus we did not place more specific hypothesis.

## Methods

This study was conducted in accordance with the Declaration of Helsinki, and it was approved by the joint Ethics Committee of the Hospital District of Southwest Finland and University of Turku (15.03.2011) §95, ETMK: 31/180/2011. Informed written consents were obtained from parents before MRI scans were conducted.

### Participants

Twenty-eight full-term born healthy infants (Table 1) were randomly recruited from the FinnBrain Birth Cohort Study (11) to be included to (f)MRI scans (functional data were scanned during year 2015). Exclusion criteria were interviewed during the recruitment phone calls and included perinatal complications of neurological involvement, less than 5 points in the 5 min Apgar, previously diagnosed central nervous system anomaly, gestational age at delivery less than 32 weeks and birth weight less than 1500 g.

**Table 1.** Demographics of subjects included in GICA and LEiDA analyses. Maternal monthly income is divided into five categories (1=<1000€ / 2=1001-2000€ / 3=2001-3000€ / 4=3001-4000€ / 5= >4000€). Maternal education was trichotomized (1 = High school graduate or lower; 2 = College degree; 3 = University degree). Frequency of maternal use of alcohol or illicit substances during pregnancy were trichotomized (more than 1-2 times a month / 1-2 times a month / less frequently). Abbreviations: M = Mean; SD = Standard deviation; MAD = Mean absolute deviation; EPDS = Edinburgh postnatal depression scale 10-point questionnaire; SCL-90 = Symptom checklist anxiety questionnaire.

|  | Variable | Whole sample<br>N=20 | Male<br>N = 11 | Female<br>N = 9 |
| --- | --- | --- | --- | --- |
|  | *M ± SD (range); For Apgar points, the median is reported* |  |  |  |
| Neonates | Age from birth (weeks) | 3.72 ± 1.11<br>(1.57 – 5.71) | 3.47 ± 1.14<br>(1.57 – 5.14) | 4.03 ± 1.06<br>(2.71 – 5.71) |
|  | Age from term (weeks) | 3.64 ± 0.71<br>(2.43 – 4.71) | 3.34 ± 0.63<br>(2.43 – 4.43) | 4.00 ± 0.65<br>(3.00 – 4.71) |
|  | Gestational age during scanning (weeks) | 43.71 ± 0.72<br>(42.43 – 44.71) | 43.42 ± 0.73<br>(42.43 – 44.57) | 44.06 ± 0.56<br>(43.29 – 44.71) |
|  | Gestational weight when born (g) | 3578 ± 343<br>(3085 – 4395) | 3536 ± 295<br>(3105 – 3980) | 3628 ± 406<br>(3085 – 4395) |
|  | Head circumference when born (cm) | 35.25 ± 1.11<br>(33.0 – 37.5) | 35.50 ± 1.12<br>(34.0 – 37.0) | 34.94 ± 1.07<br>(33.0 – 37.0) |
|  | Apgar points at 1 min (MAD) | 9.00 (1.45)<br>(3 – 10) | 9.00 (1.64)<br>(3 – 10) | 9.00 (1.21)<br>(3 – 9) |
|  | Apgar points at 5 min (MAD) | 9.00 (0.56)<br>(6 – 10) | 9.00 (0.36)<br>(8 – 10) | 9.00 (0.79)<br>(6 – 10) |
| Mothers | Maternal age (years) | 29.05 ± 4.44<br>(19 – 37) | 28.64 ± 4.74<br>(19 – 37) | 29.56 ± 4.28<br>(24 – 36) |
|  | Pre-pregnancy BMI (kg/m<sup>2</sup>) | 25.60 ± 3.53<br>(20.03 – 33.06) | 25.15 ± 3.77<br>(20.03 – 31.24) | 26.15 ± 3.35<br>(21.05 – 33.06) |
|  | EPDS score | 3.65 ± 3.30<br>(0 – 11) | 4.00 ± 2.45<br>(0 – 8) | 3.22 ± 4.37<br>(0 – 11) |
|  | SCL-90 score | 2.95 ± 3.15<br>(0 – 10) | 4.00 ± 3.31<br>(0 – 10) | 1.67 ± 2.55<br>(0 – 8) |
|  | Composite score | 6.60 ± 5.69<br>(0 – 18) | 8.00 ± 5.12<br>(0 – 17) | 4.89 ± 6.17<br>(0 – 18) |
|  | *Frequencies* |  |  |  |
|  | Maternal monthly income (€): |  |  |  |
|  | <1000 | 4 | 1 | 3 |
|  | 1001-2000 | 11 | 6 | 5 |
|  | 2001-3000 | 4 | 4 | 0 |
|  | 3001-4000 | 1 | 0 | 1 |
|  | >4000 | 0 | 0 | 0 |
|  | Maternal education level: |  |  |  |
|  | Low | 7 | 2 | 5 |
|  | Mid | 6 | 3 | 3 |
|  | High | 7 | 6 | 1 |
|  | Maternal use of alcohol during pregnancy (yes/no) | 4/16 | 2/9 | 2/7 |
|  | Frequency of maternal use of alcohol during pregnancy | 0/1/3 | 0/1/1 | 0/0/2 |
|  | Maternal use of illicit substances during pregnancy (yes/no) | 1/19 | 0/11 | 1/8 |
|  | Frequency of maternal use of illicit substances during pregnancy | 0/0/1 | 0/0/0 | 0/0/1 |

### Demographics and maternal prenatal psychological distress measures

Obstetric data was obtained from the Finnish Medical Birth Register of the National Institute for Health and Welfare (www.thl.fi). All questionnaires assessing maternal psychological health were filled in by the mothers during the 24^th^ gestational week. Maternal depressive symptoms during pregnancy were assessed by implementing the Edinburgh Postnatal Depression Scale (EPDS) and to assess maternal anxiety symptoms with the Symptom Checklist 90 (SCL-90). SCL-90 and EPDS scores were summed to generate a measure of maternal psychological distress (PSE composite score).

### MR image acquisition

For a detailed description of the visit, preparations, hearing protection, MR sequences etc. please see our prior report (12). All scans were carried out during natural sleep at the gestation corrected age of 25.85 ± 7.72 days. Each infant underwent an MRI scanning session of the brain, including a 6-minute resting-state fMRI sequence, conducted with Siemens Magnetom Verio 3T MRI scanner (Siemens Medical Solutions, Erlangen, Germany) equipped with a 12-element Head Matrix coil. As part of a maximum of 60-minute scan, we acquired 1) T1 -and T2-weighted anatomical scans at 1mm^3^ spatial resolution and 2) a 6-minute duration EPI (Echo-planar imaging) sequence with 42 slices with voxel size of 3 × 3 × 3 mm, TR 2500 ms, TE 30 ms, FOV of 216 × 216 mm and flip-angle (FA) of 80 degrees.

### Image preprocessing

The generated images of 20 subjects were processed with MELODIC toolbox of FSL: motion correction, slice timing correction, brain extraction, spatial smoothing FWHM of 5 mm, grand-mean intensity normalization of the entire 4D dataset and high-pass temporal filtering (σ = 100 s). The images were co-registered to UNC infant T2 template space with linear full search and 12 degrees of freedom (DOF) using FLIRT. Subject-level ICA was used for separating noise and signal components for manual denoising. Subject-level ICA yielded 24-45 components per subject, out of which on average 51.2 % (35.0 – 60.7 %) were classified as noise components and regressed out of the data with *fslregfilt*.

### Spatial alterations in brain networks

We first performed group ICA (GICA), which identifies networks that are common to all participants. Based on our prior work, the number of GICA components was set to 40 (12). For this data set, out of GICA runs with 30, 40, 50, 60 and automatic dimensionality estimation (yielding 111 components), setting the number of components to 40 generated a well-balanced compromise of plausible signal and noise components. We then used dual regression that uses the group level ICA maps to perform multivariate temporal regression of the individual component time courses to yield subject specific spatial maps, whose properties can then be investigated in voxel-wise general linear model (GLM) with chosen covariates.

### Temporal alterations in brain networks

To apply LEiDA we first obtained the average fMRI signals in 90 cortical and subcortical areas defined according to the UNC neonate AAL atlas (Automated Anatomical Labelling) using *fslmeants* from *FSL*. The time courses were then bandpass filtered (0.02 – 0.10 Hz), the analytic phase obtained using the Hilbert transform, and the leading eigenvectors of the phase coherence matrices were calculated at each time point (10). The eigenvectors were then clustered via K-means clustering based on cosine similarity and 200 replicates. For the purposes of this explorative study, the number of clusters (K) was varied between 2 and 8, based on previous studies indicating that the optimal number of intrinsic networks is typically between 3 and 8. We placed an emphasis on describing the brain networks that emerge from each clustering solution and exploring their associations to PSE (rather than formally assessing the best clustering solution). Each clustering yielded K patterns of phase-relationships in brain activity - or Functional Connectivity (FC) states - whose probability of occurrence can be used in statistical inference (10).

For the purposes of this study 20 out of 27 scanned neonates had usable data, i.e., 6 subjects were excluded due to clear over motion and one participant due to unsuccessful K means clustering result. Mean motion parameters of the included subjects were 1.039 mm and 0.349 mm for mean and relative displacements, respectively, as reported in FSL’s MCFLIRT.

### Statistical analyses

Brain network metrics were the main dependent variable, and the composite PSE score was the independent variable of interest. We used FSL randomize to test associations between GICA maps and JASP (version 0.16.1.0) to test associations of LEiDA-derived network probabilities of occurrence with ANCOVA models, including Levene’s test and normality checks. We included as covariates / controlled for: neonate sex, gestational age at scanning, and age from birth in all models. In sensitivity analyses we additionally controlled for neonate birth weight and maternal pre-pregnancy Body Mass Index (BMI). We corrected the statistical tests for multiple comparison with Bonferroni correction over the number of clusters, i.e., for each independent hypothesis tested.

## Results

### Spatial alterations in brain networks

GICA networks were expectedly well aligned with our prior report (12). We did not find associations between GICA defined networks and PSE using dual regression.

### Temporal alterations in brain networks

The repertoire of network patterns – or FC states - captured with LEiDA was found to reveal canonical resting-state networks described in the literature (Supplemental figure 1) replicating prior reports with static and dynamic methods (12,13). While the patterns obtained when clustering with K=2 to K=5 did not show statistically significant associations with PSE, when clustering the patterns into six clusters, we observed a robust effect of maternal distress composite score on the probability of occurrence of the frontoparietal network (Figure 1, Supplemental figure 2) after controlling for neonate sex, gestational age at scanning and age from birth, F (1, 18) = 15.774, p = 0. 001, ω^2^ = 0.293, which survived Bonferroni correction over the number of independent hypotheses tested (corrected p = 0.006). This association was specific to the implicated frontoparietal network (Supplemental figure 3).

**Figure 1.**
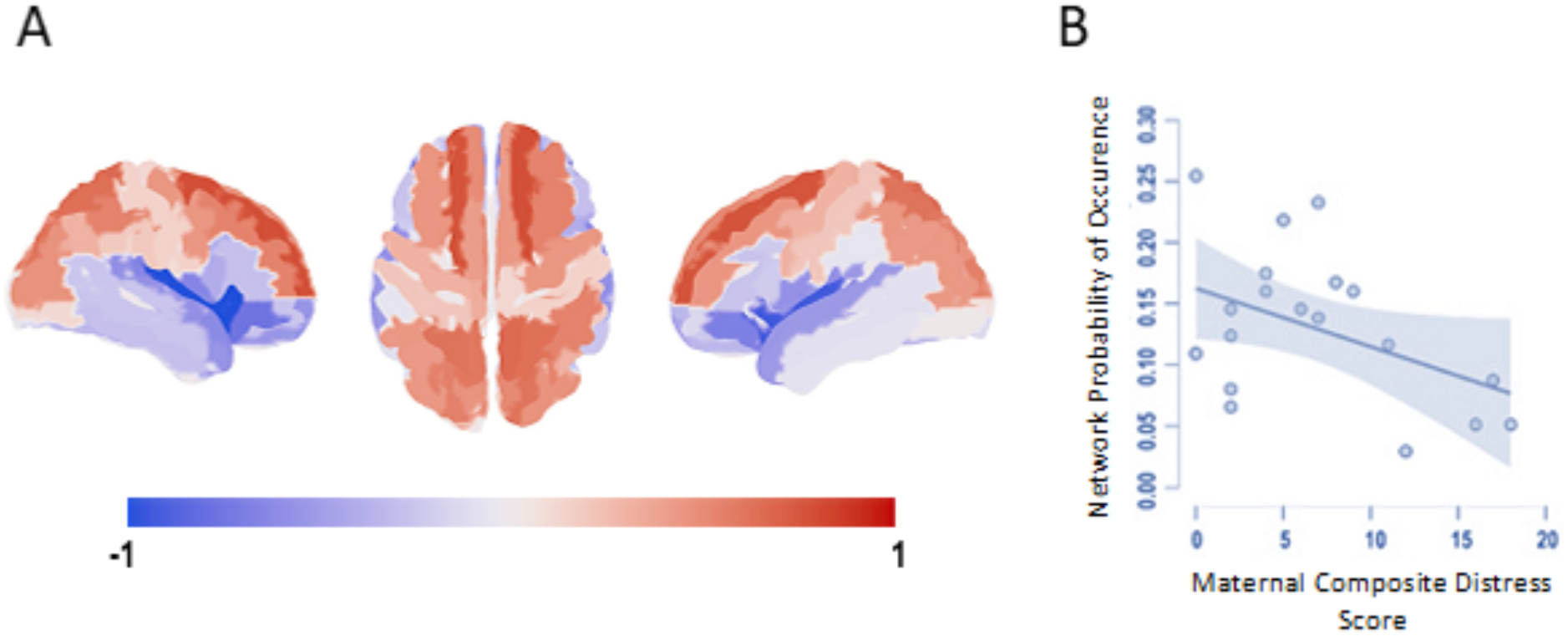
Maternal prenatal distress associates negative with the probability of occurrence of a frontoparietal network: A) visualization of the brain network #3 identified from clustering solution with 6 clusters; B) scatter plot of the association between probability of occurrence and PSE (Spearman’s rho = -0.716, p=0.001, Bonferroni-corrected p = 0.006).

### Sensitivity analyses

Levene’s test and normality checks were carried out and the assumptions met. The effect sizes were smaller but statistically significant in our sensitivity analyses where maternal-pre-pregnancy BMI and neonate birth weight were additionally controlled for, F (1, 18) = 7.757, p = 0.015, ω^2^ = 0.170). This network also appeared in clustering solutions with seven (FC state 4) and eight (FC state 4) networks and had similar associations with PSE composite score in fully controlled ANCOVA models that included all covariates of the sensitivity analyses (K = 7, F [1, 18] = 7.997, p = 0.014, ω^2^ = 0.183; K = 8, F [1, 18] = 5.637, p = 0.034, ω^2^ = 0.127). This gives assurance that the effects are not subject to clustering solutions.

## Discussion

The current study probed the links between PSE and neonatal whole brain networks for both spatial and dynamic features. We found a statistically robust negative association with the probability of occurrence (stability) of a frontoparietal network and PSE that was defined as a composite score of depressive and anxiety symptoms. No associations were found between static network FC and PSE. To bring our findings to context, we briefly review the four prior fMRI studies that have addressed the links between PSE and neonate brain function.

Scheinost et al. (5) used two data sets / cohorts focusing on very/extremely preterm neonates with or without PSE, defined as a binary classification for maternal prenatal diagnosis of depression, compared with term controls. Seed-based connectivity analyses revealed that very preterm subjects exhibit reduced amygdala connectivity to subcortical structures when compared to term controls. Extremely preterm neonates with PSE have further reduced amygdala FC to subcortical structures when compared to extremely preterm neonates without PSE indicating that PSE further reduces amygdala FC to subcortical structures.

Posner et al. (6) studied 64 infants with (n=20) and without (n=44) in utero exposure to prenatal maternal depression. PSE-exposed infants had more negative functional connectivity between amygdala and dorsal prefrontal cortex (AMG-dPFC) and altered AMG-dPFC FC further associated with greater prenatally assessed stress responses (fetal heart rate measurements). Of note, this study also assessed structural connectivity and found decreased structural connectivity between right amygdala and right ventral prefrontal cortex in neonates with PSE exposure.

Qiu et al. (7) studied older infants, 24 infants at 6 months age (12 males and 12 females). PSE was assessed via self reports with Edinburgh prenatal depression scale (EPDS) at 26 weeks of gestation and 3 months after delivery. Pre- and postnatal EPDS scores that were found to be similar. They found a positive association between EPDS scores and infant left amygdala FC between left temporal cortex, insula and bilateral anterior cingulate cortex (ACC), medial prefrontal cortex (mPFC) and ventromedial prefrontal cortex (vmPFC).

Another study from the same cohort studied similar associations in older children. Soe et al. (8) scanned 128 children at 4.4-4.8 years of age. PSE was measured with maternal self reports via EPDS-questionnaire at 26 wk of gestation, 3 months, 1, 2, 3 and 4.5 years after delivery. The authors used prenatal EPDS scores, a mean EPDS score value of postnatal EPDS scores and their difference as variables of interest. They found no associations in boys. In girls they found that prenatal PSE (depressive symptoms) associated with FC of amygdala and cortico-striatal circuitry (orbitofrontal cortex, insula, anterior cingulate cortex, temporal pole, striatum). Further, in girls only, greater pre-than post-natal depressive symptoms associated with lower FC between left amygdala and bilateral anterior cingulate cortex, left caudate; right amygdala and left orbitofrontal cortex, insula and temporal pole.

Taken together, the main method for studying links between PSE in newborns has been seed-based connectivity of the amygdala, and implicated regions encompass striatal and frontal brain areas with PSE associating to decreased connectivity. All prior studies, including our own, is cross sectional, and definition of maternal prenatal “stress” is variable across studies. Our study used a general population-based, non-clinical sample and used a continuum of symptom scores similar to some prior reports (7,8). Although no clear clinical cut-off scores have been established for EPDS and SCL, generally a score of 10 has been implicated as a clinically meaningful threshold for symptoms of depression or anxiety in pregnancy (11). It is important to note that maternal prenatal stress is only partly captured by depressive and anxiety measurements. More comprehensive approaches may reveal patterns of stress exposure that better explain offspring outcomes at peripartum and postnatally (14).

During normal development, amygdala seed-based FC becomes negatively associated with age at many overlapping posterior brain regions found in the frontoparietal areas (15). The developmental trajectories of the connectivity indicate that frontoparietal networks are slow to achieve adult-like topology (16), which may render them vulnerable to stressors (17). Overall, there is cumulating evidence pointing towards stress exposure associated with accelerated brain maturation (16,17), and this may well be reflected in amygdalar FC to frontal and parietal regions.

The overall implication of destabilized frontoparietal network dynamics is intriguing as similar network patterns are frequently linked to later child outcomes - albeit with different fMRI methods. Frontoparietal regions have been linked to PSE in fetal studies (17) and also to behavioral phenotypes in 1-month old infants (18). Later in development, frontoparietal networks are often referred to as attention networks and are implicated in neurodevelopmental disorders (19) and mental health (20).

## Conclusions

We found that stability of a frontoparietal network is negatively associated with maternal mid-pregnancy PSE (defined as a composite score of depressive and anxiety symptoms). This study adds to the cumulating evidence that frontal brain areas and associated networks are potentially especially sensitive to PSE. There is a dire need for studies replicating and extending prior studies in larger sample sizes to define the clinical relevance of these findings.

## Acknowledgements

We thank all FinnBrain families that took part in the MRI studies, our radiographer Krisse Kuvaja for performing the imaging, and the FinnBrain Birth Cohort study staff.

## Data availability statement

The Finnish law and ethical permissions do not allow the sharing of the data used in this study.

---

**Supplemental figure 1.**
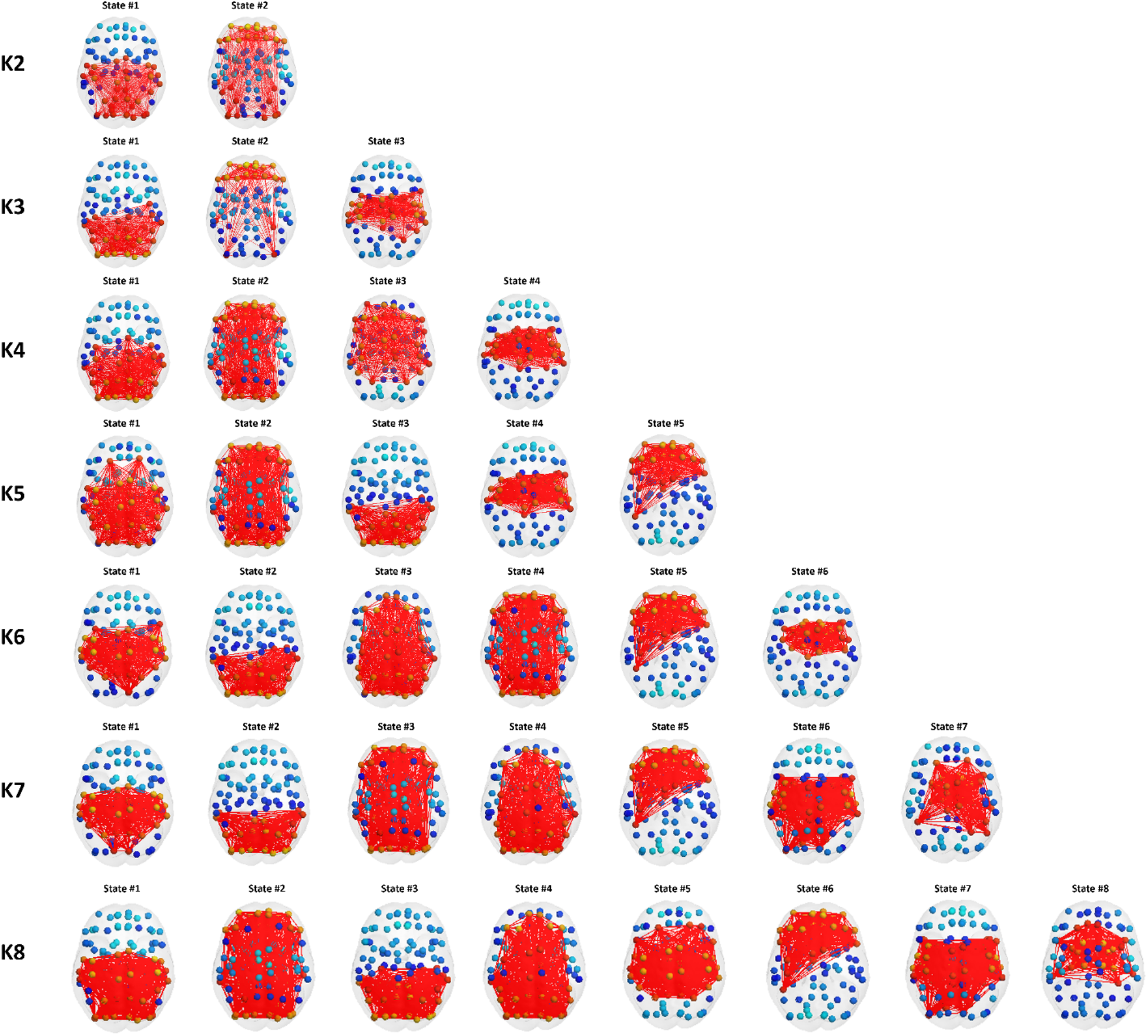
LEiDA network pyramid depicting functional brain states detected for increasing number of clusters K (with K = 2,…,8) and sorted from left to right according to probability of occurrence. Our most significant result (obtained for K=6, FC state 3) has a similar configuration through clustering solutions of K = 6 to 8 (K = 7,8 = FC state 4).

**Supplemental figure 2.**
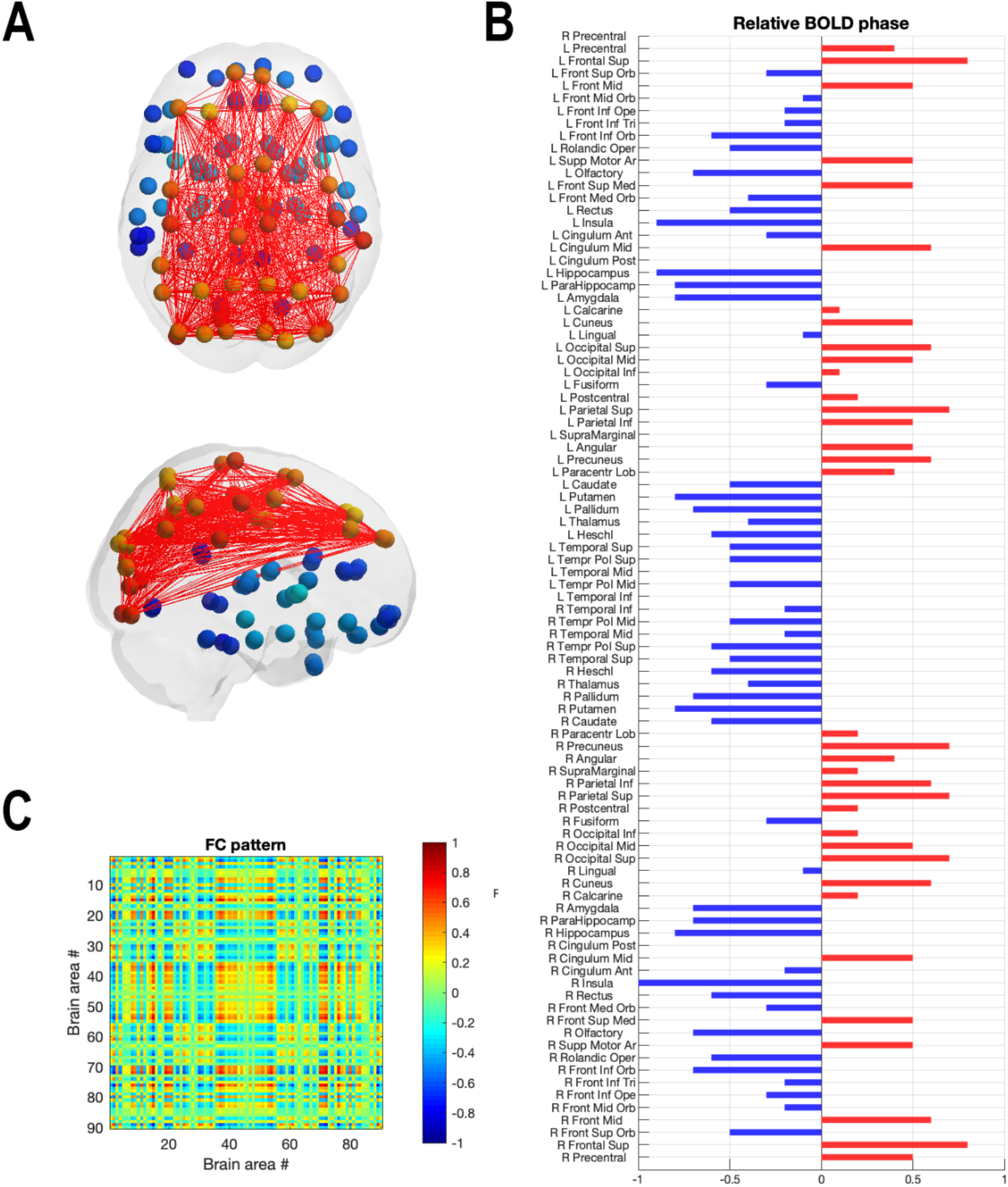
The frontoparietal network (brain state 3 when K=6): A) glass brain visualization of the network, B) the connectivity pattern in AAL atlas (90×90 brain areas), C) visualization of the brain areas that constitute the network (in red colour).

**Supplemental Figure 3.**
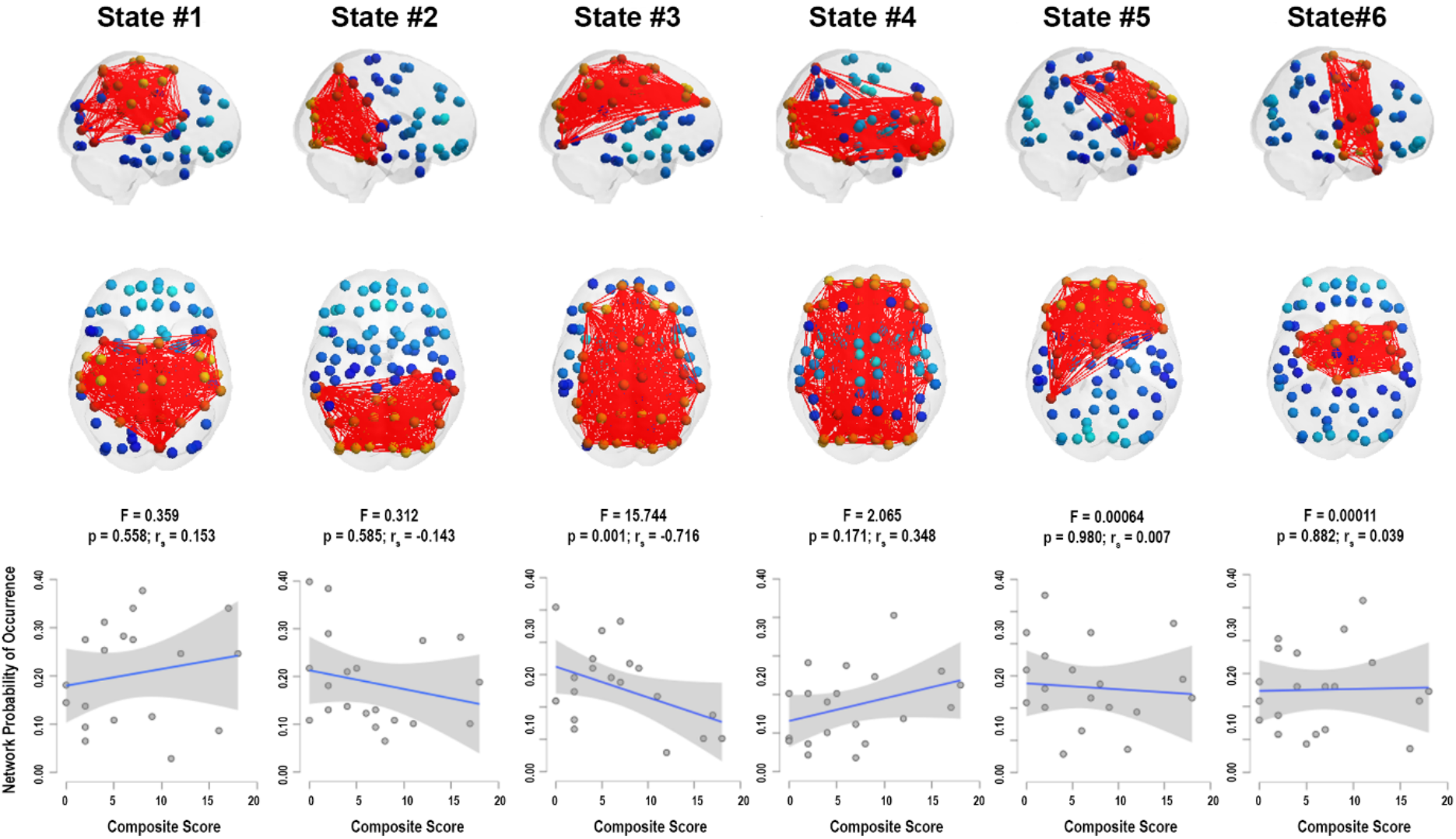
The scatter plots of the associations between the probability of occurrence across the brain states in clustering solution with six clusters. While the scatter plots show many possible associations, after controlling for neonate sex, gestational age at scanning and age from birth, most lose statistical significance (see effect size estimates and p values below the plots for Spearman partial correlations [r_s_]). The x axis’ ‘Composite score’ corresponds to the PSE composite score (generated by summation of maternal EPDS and SCL-90 questionnaire points).

